# State dependent constraints on claustrocortical communication and function

**DOI:** 10.1101/2023.09.15.557951

**Authors:** Brian A. Marriott, Alison D. Do, Coline Portet, Flora Thellier, Romain Goutagny, Jesse Jackson

## Abstract

Neural activity in the claustrum has been associated with a range of vigilance states, yet the activity patterns and efficacy of synaptic communication of identified claustrum neurons have not been thoroughly determined. Here we show that claustrum neurons projecting to the retrosplenial cortex were most active during synchronized cortical states such as non-rapid-eye-movement (NREM) sleep and were suppressed during increased cortical desynchronization in arousal, movement, and REM sleep. The efficacy of claustrocortical signaling was also increased during NREM and diminished during movement due in part to increased cholinergic tone. Finally, claustrum activation during NREM sleep enhanced memory consolidation through the phase-resetting of cortical delta waves. Therefore, claustrocortical communication is constrained to function most effectively during cognitive processes associated with synchronized cortical states, such as memory consolidation.

**Key Points:**

1. Claustrum neurons are suppressed/activated during desynchronized/synchronized cortical states
2. Claustrocortical connectivity is increased in non-REM and suppressed during movement
3. Claustrocortical communication is heightened during low cholinergic tone
4. Claustrum – modulated cortical delta waves enhance the consolidation of a labile memory

## Introduction

The claustrum provides a diffuse excitatory connection to almost all cortical regions ^1–5^. The function of the claustrum has remained somewhat enigmatic as it has been implicated in cortical synchrony, sleep, stress, memory, attention, and decision making ^6–11^. One challenge has been relating neural activity in the claustrum to these distinct processes, as the real-time measurement of identified claustrum neurons has only been reported in a select few instances ^6,7,11–16^. It is therefore unclear how these seemingly different behavioral states are supported by activity in the claustrum.

Insight into the function of neural circuits can be gained by understanding how neural activity relates to changes in brain or behavioral state ^17–20^. Across sleep-wake states, cortical networks display continuous and spontaneous changes in network activity ^20–23^. Distinct network patterns, measured by local field potential (LFP) or electroencephalographic (EEG) recordings ^23–25^, reflect different states which may or may not be accompanied by changes in behavioral output ^20,21,24^. States of cortical desynchronization, defined by prominent hippocampal theta (7-10Hz) and cortical gamma frequency (30-100Hz) and low delta frequency (0.5-4Hz) activity, are a hallmark of wake-related behaviors such as exploration, memory encoding, attention ^26–29^, and rapid eye movement (REM) sleep ^17,22^. On the contrary, synchronized cortical states are characterized by high delta power and arise during rest and non-REM (NREM) sleep and participate in processes such as memory consolidation ^21,30–32^, metabolic homeostasis ^33^, or synaptic scaling ^34,35^. Therefore, deciphering the brain state-dependent organization of neural activity informs the functional constraints placed on a population of neurons ^36^.

Recent reports in different species have described increased activity of claustrum neurons in NREM sleep ^7,13,16^ i.e. in synchronized cortical states. However, the claustrum is also proposed to function in REM sleep and states of high attentional demand such as memory encoding or decision making ^8,10,11,37,38^, suggesting activation during cortical desynchronization. This work raises the question of whether there is an optimal brain or behavioral state for claustrum communication with downstream targets in the mammalian brain. Our goal in this study is to measure how claustrum activity and the efficacy of communication with the cortex changes across different brain and behavioral states. The determination of when claustrocortical neurons effectively communicate with target regions will help elucidate how this circuit participates in state-dependent cognitive processes.

## Results

### Imaging claustrum axons across brain and behavioral states

The claustrum projection to the retrosplenial cortex has been well studied at the cellular and circuit connectivity level ^15,39–42^. However, very little is known about the activity patterns of these neurons. To measure claustrum activity in this circuit, we performed in vivo two-photon calcium imaging of claustrum axons innervating the retrosplenial cortex, in head-fixed male and female mice. Specific targeting of these neurons was achieved using a dual virus approach where AAVretro-syn-cre was injected into the retrosplenial cortex and the cre dependent calcium sensor AAV1-FLEX-jGCaMP8m ^43^ injected into the claustrum (**Figure 1A, see Methods and Materials**). Since brain regions neighboring the claustrum do not project considerably to the retrosplenial cortex ^2,15,39^ this strategy provides a claustrum specific expression of GCaMP. Two photon, thin-skull ^44^ axon imaging (**Figure 1B**) was conducted in a ‘sleep tube’ or on a self-propelled treadmill ^45^ to promote locomotion. Local field potentials were recorded near the imaging window and together with pupil size and face motion enabled the detection of quiet wake, NREM, and REM sleep while in the sleep tube ^46–48^ (**Figure 1A-E)** or periods of locomotion and quiet wakefulness on a treadmill (**Figure 1F-G**, **Methods and Materials**). Imaging was performed primarily in layers 1-3 (30-150µm from pia), and retrograde tracing showed that claustrum neurons innervating the retrosplenial cortex often project to both superficial layers (1-3) and deep-layers (5-6) (**Figure S1A-C**). Therefore, imaging superficial layers provides knowledge of the claustrum-retrosplenial input as a whole.

**Figure 1.**
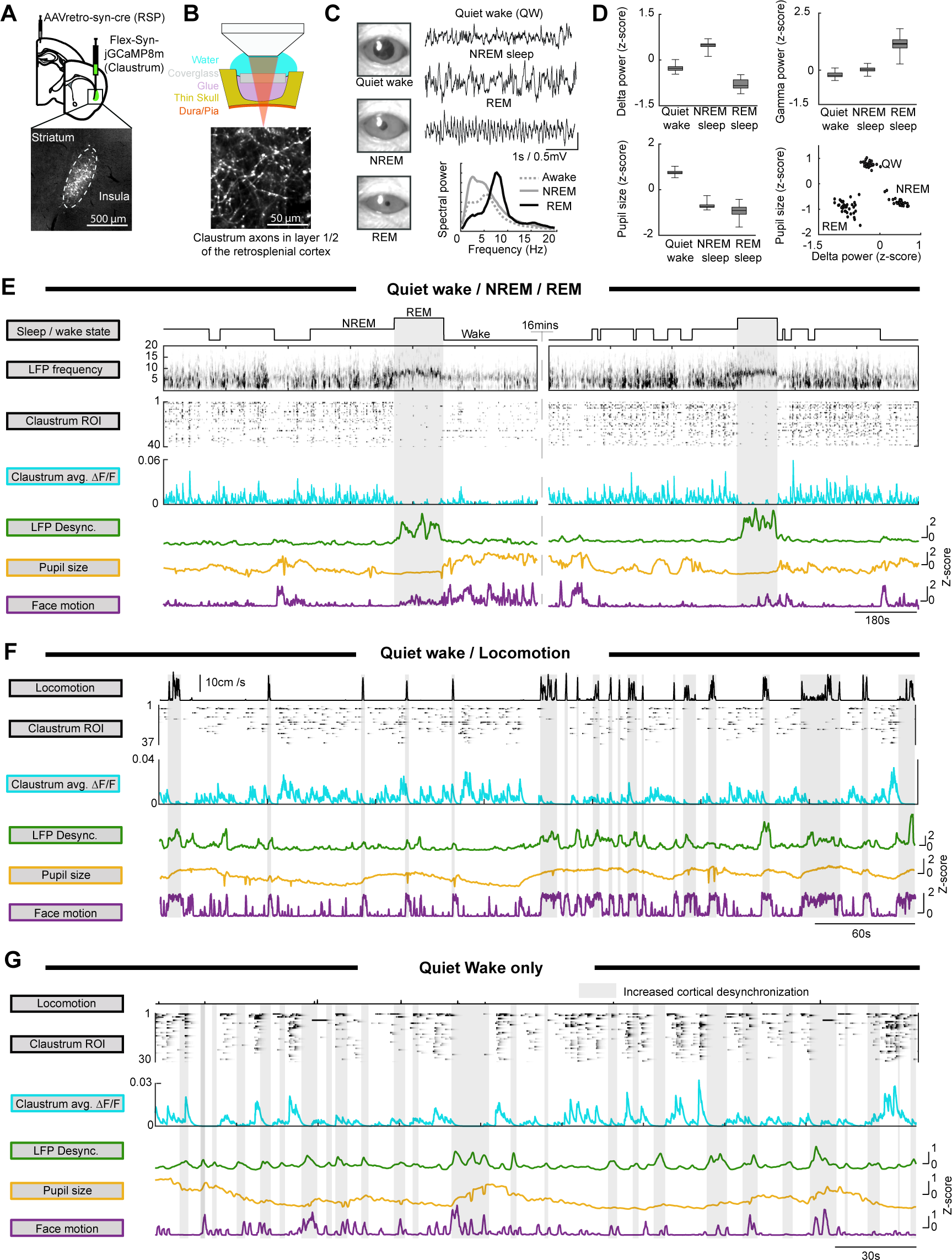
Imaging claustrum axons across brain and behavioral states. **A**: The dual virus approach, enabling the expression of GCaMP8m in claustrum neurons projecting to the retrosplenial cortex. **B**: Schematic depicting thin-skull two-photon calcium imaging. Below is an example maximum projection image from one field of view showing claustrum axon projecting to the retrosplenial cortex. **C**: Left: Pupil images during quiet wake, NREM sleep, and REM sleep. Right: Example local field potential (LFP) recordings (top) and power spectrum analysis (bottom) during sleep-wake states. **D**: Sleep-wake states are defined by specific pupil size, gamma frequency power, delta frequency power. **E**: An example session during sleep-wake transitions, showing (top to bottom): The spectrogram of the LFP, example claustrum ROIs, the mean claustrum population activity, the LFP desynchronization (gamma/delta power ratio), pupil size, and face motion. **F**: Similar to E, with a mouse transitioning between quiet wakefulness and locomotion on a self-paced treadmill. **G**: The same as F but focused on a period of quiet wakefulness. Grey shading indicates REM (**E**), locomotion (**F**), or spontaneous increases in cortical desynchronization (**G**).

### Claustrum-retrosplenial activity is suppressed during REM

It is currently unclear how projection defined claustrum neural activity is organized across sleep-wake states ^7,13,37,49^. Therefore, we measured and compared claustrum activity as mice cycled between NREM, REM, and wake. For each session, we measured the changes in fluorescence of individual axon regions of interest (ROI) within a field of view or the averaged ROI activity (claustrum population activity). Every session (n = 35) in every mouse (n = 6) showed decreased claustrum population activity during REM relative to NREM (**Figure 2A-B, FigureS1D-I**), and 34/35 sessions showed decreased activity in REM relative to wake. The majority (62%) of individual ROI were silenced during REM and 71% were preferentially active during NREM sleep (**Figure 2C**). The 10% of ROI that were preferentially active during REM had a lower overall activity rate than the population of NREM preferring ROIs (**Figure S1D-F**). Activity during quiet wakefulness was consistently lower than NREM sleep with only 19% of ROIs being preferentially active during wake. Therefore, claustrum neurons projecting to the retrosplenial cortex are preferentially activated during NREM and are predominantly inhibited during REM-associated desynchronization.

**Figure 2.**
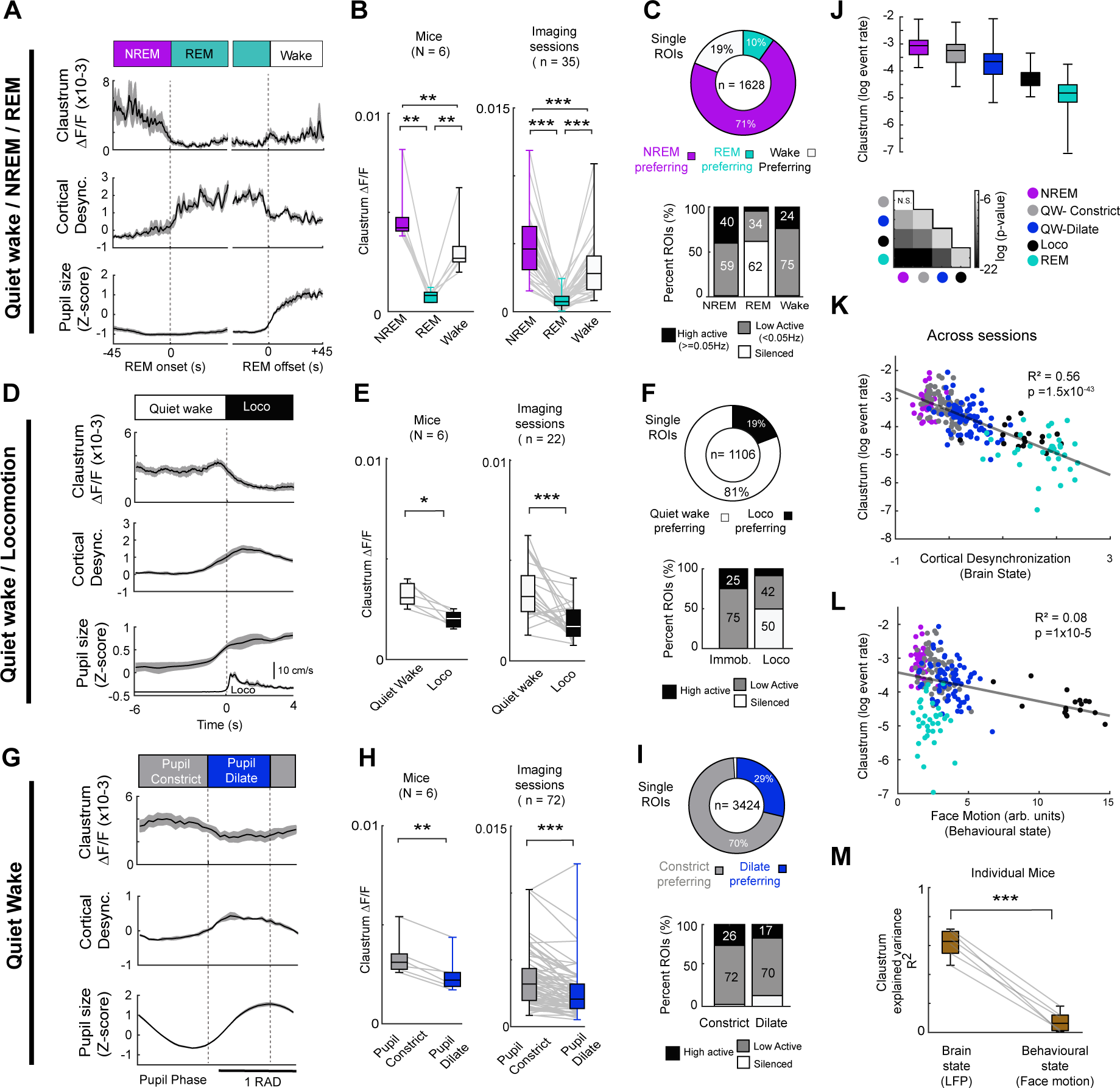
Claustrum activity scales as a function of brain state. **A**: The time-course of claustrum activity during NREM – REM - wake transitions (mean ±s.e.m., N = 6). **B**: Data averaged across mice (left) and sessions (right), showing differences between NREM and REM (Mice: t_(5)_ = 6.7, p = 0.002; Sessions: t_(34)_ = 11.1, p=1×10^-12^), REM and wake (Mice: t_(5)_ = 4.3, p = 0.007; Sessions: t_(34)_ = 6.9, p=5×10^-8^), and Wake and NREM (Mice: t_(5)_ = 9.4, p = 0.001; Sessions: t_(34)_ = 11.9, p = 3×10^-13^). **C**: The single ROIs analysis showing the proportion of ROI with higher event rates in NREM, REM, or wake. Below is the breakdown of activity in each state, sorted into silent, low active, and high active. **D-F**: The same as A-C but for sessions on the treadmill. Locomotion is shown in the inset. Activity during locomotion was reduced across mice (t_(5)_ = 3.9, p = 0.01) and sessions (t_(21)_ = 4.5 p = 1×10^-4^). **G-I**: The same as **D-F** but for periods of quiet wakefulness from sessions on both the treadmill and sleep tube. Activity was reduced during pupil dilation across mice (t_(5)_ = 6.5, p = 0.001) and sessions (t_(71)_ = 6.6, p = 6×10^-9^). **J**: The overall Ca^2+^ event rate for claustrum ROIs across brain states in each session. **K**: The relationship between the mean ROI event rate within a given brain state versus cortical desynchronization for each session. State color is as indicated in J. **L**: The same as **K** but for face motion as an index of behavioral state. **M**: The R^2^ between claustrum activity and brain state or behavioral state for each mouse (t_(5)_ = 13.7, p= 3.7×10^-5^). Box plots throughout show median, interquartile range, and maximum/minimum values. *p<0.05, **p<0.01, ***p<0.001.

### Claustrum-retrosplenial activity is reduced during locomotion and arousal

Following several sessions on the sleep-tube, mice were trained to run on a self-paced treadmill and claustrum activity was compared between periods of quiet wake and locomotion (**Figure 1F**, **Figure 2D**, **Figure S2**). Claustrum activity was suppressed during locomotion in all mice and in the majority of sessions (**Figure 2E**). Most axon ROIs showed higher activity event rates during quiet wake relative to locomotion (81% preferred quiet wake) with half the ROI not exhibiting any calcium events during locomotion (**Figure 2F**). During quiet wakefulness, mice exhibit spontaneous fluctuations in brain state where changes in pupil dilation/constriction signal increases/decreases in arousal respectively ^47,48,50,51^. Therefore, we measured changes in claustrum activity as a function of pupil diameter during quiet wake on either the sleep tube or treadmill (**Methods and Materials**). Across imaging sessions and mice, claustrum activity was suppressed during dilation (increased arousal) and predominantly active during pupil constriction (decreased arousal, **Figure 2G-I**). The majority of ROI (70%) were more active during constriction than dilation indicating that the activity of claustrum neurons is reduced during spontaneous increases in arousal. The correlation coefficient between each ROI and pupil diameter showed a negative overall correlation across both wake and NREM sleep (**Figure S3A-L**). Collectively, these data show that the claustrum-retrosplenial pathway is more active during states associated with synchronized cortical activity (NREM and low arousal) and less active during states associated with cortical desynchronization (locomotion, increased arousal, and REM sleep), with claustrum activity undergoing a ∼5-fold change between NREM and REM sleep (**Figure 2J**).

Small changes in behavioral state influence neural activity across brain regions ^51–53^. However, claustrum activity appears to be related to brain state changes rather than changes in behavioral state, as REM and locomotion have similar brain state desynchronization and the lowest claustrum activity yet opposing behavioral outputs. To test this hypothesis, we compared how claustrum activity scaled as a function of brain state versus behavioral state. Brain state was measured as the ratio between gamma power to delta power (cortical desynchrony). Facial movement of the vibrissae occurs throughout wake-sleep states and therefore, we used this face motion as a proxy for behavioral state similar to previous reports ^52,54^. We found a strong relationship between cortical desynchrony and claustrum activity (R^2^=0.56, p<0.001, **Figure 2K**) which was also present within individual sessions (**Figure S3M-N**). Behavioral state did not strongly covary with claustrum activity across sessions (R^2^=0.08, **Figure 2L**). Within individual sessions, face motion was negatively correlated with claustrum population activity and individual ROI (**Figures S3C-L**). However, the partial correlation between claustrum activity and facial motion (taking into consideration cortical desynchronization), was reduced (**Figure S3M-N**) suggesting that the relationship between claustrum activity and face motion was mostly driven by the correlation between face motion and brain state. Therefore, claustrum activity was more correlated with changes in brain state than with behavioral state (**Figure 2M**).

### Claustrocortical communication is increased during NREM sleep and reduced during locomotion

Quiet wakefulness and NREM sleep - two behavioral states characterized by the presence of slow-wave cortical activity - had the highest claustrum activity levels. We therefore determined the correlation coefficient between claustrum population activity and the LFP amplitude fluctuations across all frequencies ranging from 0.5 to 100Hz. The delta frequency band had the highest positive correlation with claustrum activity across all mice (**Figure 3A-D**). We compared the correlation between delta power and claustrum population activity (**Figure S3D-J**) in the two states. The time-lag at maximum correlation underwent a positive shift between wake and NREM indicating that claustrum activity was more correlated with increases in future delta frequency amplitude during NREM relative to wake (**Figure 3C**). Therefore, claustrum activity is more likely to participate in shaping the dynamics of ongoing delta wave fluctuations during NREM, whereas the cortex may be responsible for shaping claustrum activity during wakefulness.

**Figure 3:**
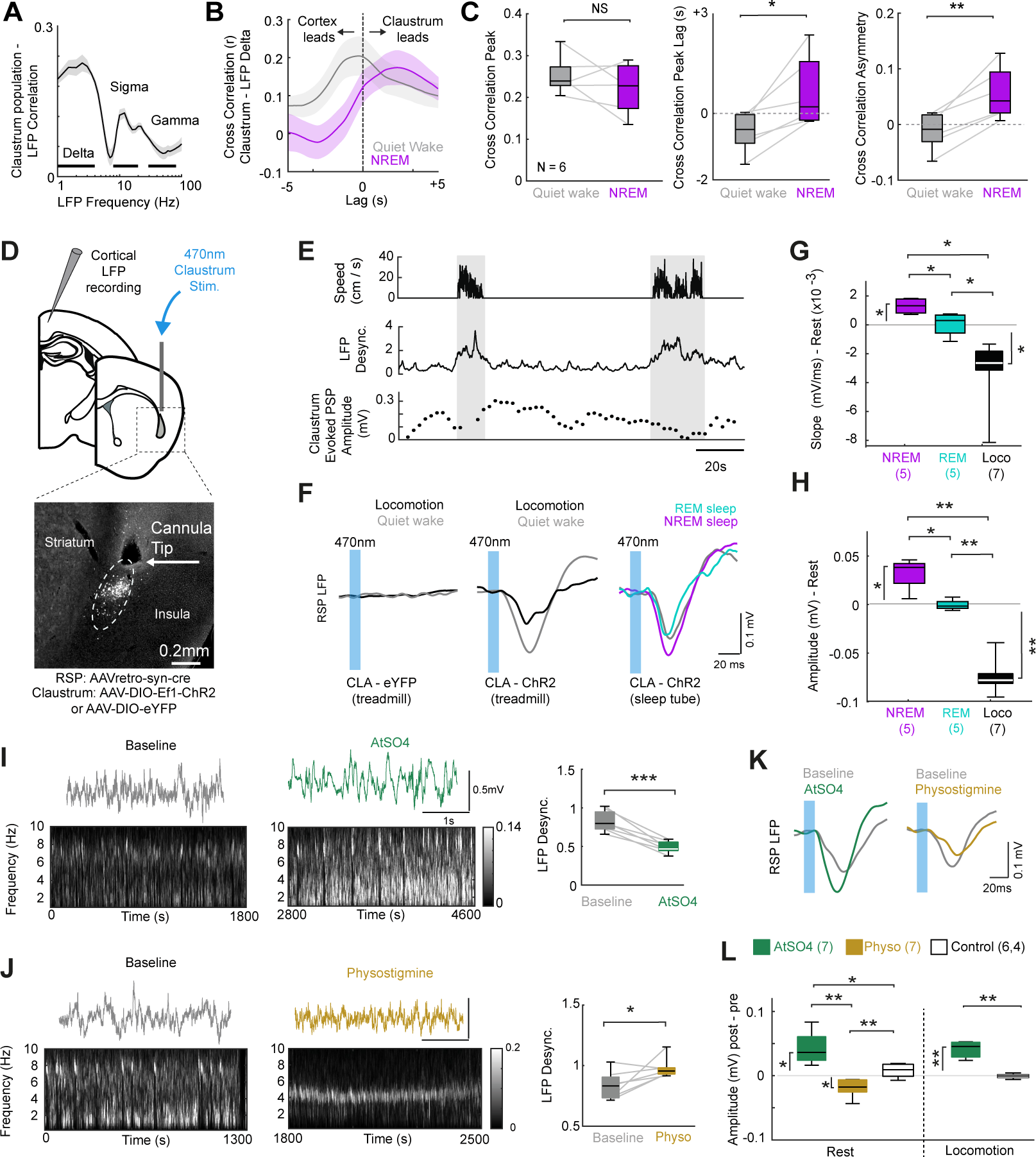
Claustrocortical signaling is most effective during NREM sleep and reduced by high cholinergic tone and movement. **A**: The mean ± s.e.m. correlation between claustrum population activity and LFP frequency bands. **B**: The mean time-lagged cross correlation between claustrum activity and cortical delta power for one mouse. **C**: The peak cross correlation for all mice (left, t_(5)_ = 1.13, p = 0.31), time-lag at maximum correlation (middle, t_(5)_ = 3.64, p = 0.015), and cross correlation asymmetry (right, t_(5)_ = 5.45, p = 0.003). **D**: Top: Local field potentials were recorded while optogenetically activating claustrum-retrosplenial neurons. Bottom: ChR2 expression and the location of the optic fibre in the claustrum. **E**: Example experiment showing the claustrum ePSP response during rest and locomotion. Each point reflects moving average of five pulses. **F**: The averaged ePSPs following 10ms light pulses for representative sessions in control (eYFP, left panel), or ChR2 mice (middle and right panels). **G**: Group mean differences in ePSP slope between NREM and rest (t_(4)_ = 4.96, p = 0.031), NREM and REM (t_(4)_ = 7.33, p = 0.011) REM and locomotion (t_(10)_ = 2.8, p = 0.04), NREM and locomotion (t_(10)_ = 10.48, p = 6.2×10^-6^) and rest and locomotion (t_(6)_ = 10.72, p = 1×10^-4^), normalized to quiet wakefulness. Animal numbers are indicated below. **H**: Group mean differences in fPSP amplitude between NREM and rest (t_(4)_ = 4.58, p = 0.026), NREM and REM (t_(4)_ = 4.77, p = 0.026), REM and locomotion (t_(10)_ = 8.27, p = 4.4×10^-5^), NREM and locomotion (t_(10)_ = 10.48, p = 6×10^-6^), and locomotion and rest (t_(6)_ = 10.73, p=1×10^-4^). **I**: Example spectrograms of retrosplenial LFPs before and after administration of atropine sulfate (10mg/kg), and the change in cortical desynchronization (baseline = 0.82±0.05, atropine= 0.49±0.03; t_(6)_ = 7.1, p = 4×10^-4^). **J**: The same as **F** but following physostigmine (baseline = 0.83±0.04, physostigmine = 0.98±0.03; t_(6)_ = 2.54, p = 0.04). **K**: Example ePSPs in the retrosplenial cortex following atropine (left panel) or physostigmine (right panel). **L**: Group data normalized to baseline showing atropine increased the amplitude of the ePSP (t_(6)_ = 4.61, p = 0.015) and physostigmine decreased ePSP during rest (t_(6)_ = 3.99, p = 0.016). Both were significantly different from the response following control injections of saline (atropine: t_(12)_ = 3.4, p = 0.016, physostigmine: t_(12)_ = 4.41, p = 0.004). Atropine increased the ePSP amplitude during locomotion relative to baseline (t_(5)_ = 8.19, p = 0.001) and relative to saline (t_(8)_ = 6.45, p = 0.002). Mice did not consistently exhibit locomotion following physostigmine so no ePSP data is reported here. *p<0.05, **p<0.01, *** p <0.001.

Changes in behavioral state also impact the efficacy of synaptic transmission ^55,56^ and sensory responses in the cortex ^22,51,57–60^. Previously it was noted that claustrum activation is less effective during active wake ^7,41,45^, but this has not been systematically measured. To study the strength of communication between the claustrum and cortex, we expressed the excitatory Channelrhodopsin^61^ (ChR2 or ChETA) in claustrum-retrosplenial projections and optically stimulated these neurons while measuring the extracellular evoked post-synaptic potential (ePSP) in the retrosplenial cortex (**Figure 3D**). The slope and amplitude of the ePSP was larger in NREM than REM and reduced during locomotion relative to rest (**Figure 3E-G**). There was a negative correlation between the size of the ePSP and both cortical desynchrony and face motion within wake and sleep sessions (**Figure S4A-C**). The stimulation efficacy was reduced during increased cortical desynchrony and facial movement, even in the absence of locomotion. Claustrum stimulation did not impact running speed (**Figure S4D-E**). This suggests that both brain and behavioural state impact communication at claustrocortical synapses.

### High muscarinic cholinergic tone suppresses claustrocortical communication

It is known that neuromodulators such as acetylcholine participate in the generation of desynchronized brain states ^19,50,62,63^. For example, forebrain cholinergic activity is maximal during exploration and movement, and minimized during NREM sleep ^54,64–66^. We investigated whether the observed state-dependent variability in signaling from the claustrum to the retrosplenial cortex could be mediated by acetylcholine. We blocked muscarinic receptors using atropine sulfate (10mg/kg), which led to a decrease in cortical desynchronization and the emergence of large cortical slow waves (**Figure 3F**) as previously reported ^19,20,56,67,68^. The amplitude of claustrum evoked fPSPs in the retrosplenial cortex was increased during rest and locomotion compared to baseline following atropine (**Figure 3F-I**). Inversely, enhancing cholinergic tone with physostigmine increased cortical desynchronization and reduced the amplitude of claustrum evoked ePSPs (**Figure 3G-I**). This suggests muscarinic cholinergic signaling can attenuate communication between claustrum neurons and the cortex similar to the suppression of excitatory synaptic transmission in thalamocortical and corticocortical circuits ^69–71^. As cholinergic neurons innervate both the claustrum ^39,72^ and the cortex ^73,74^ the effects of acetylcholine on the signaling between claustrum and cortex could arise at the level of the claustrum or at claustrocortical synapses. Indeed, muscarinic receptor blockage also increased the spontaneous activity of claustrum neurons projecting to the retrosplenial cortex (**Figure S5**). This result suggests that acetylcholine could be responsible for the claustrum inhibition associated with cortical desynchronization but the cellular mechanisms underlying cholinergic control of claustrocortical communication require further investigation.

### Claustrum modulated delta waves support memory consolidation

Claustrum neurons were most active and evoked the largest post-synaptic response during NREM (**Figures 1-3**). Therefore, we measured the effect of low frequency claustrum stimulation on cortical network activity during NREM. As reported previously ^7^, claustrum stimulation could induce delta waves in the cortex, with a structure similar to spontaneous delta waves (**Figure 4A-B**). We also found a decrease in gamma frequency activity following the light pulse, consistent with inverse relationship between delta and gamma frequency power (**Figure 4C**). However, claustrum stimulation was not always effective at producing a delta wave. The phase of ongoing low frequency activity had an important impact on the ability to induce a delta wave. Stimulation during falling phase failed to elicit a delta wave and stimulation on the rising phase elicited a delta wave on 39±6% of the trials, significantly above that expected by the spontaneous occurrence of delta waves (28±2% of low frequency events, **Figure 4D, Methods and Materials**). Therefore, the impact of claustrum activation on the cortex depends on brain state and the phase of ongoing cortical activity in the milliseconds prior to stimulation. These results suggest that rather than driving delta frequency cortical activity, claustrum-retrosplenial circuits modify the timing of cortical delta waves.

**Figure 4.**
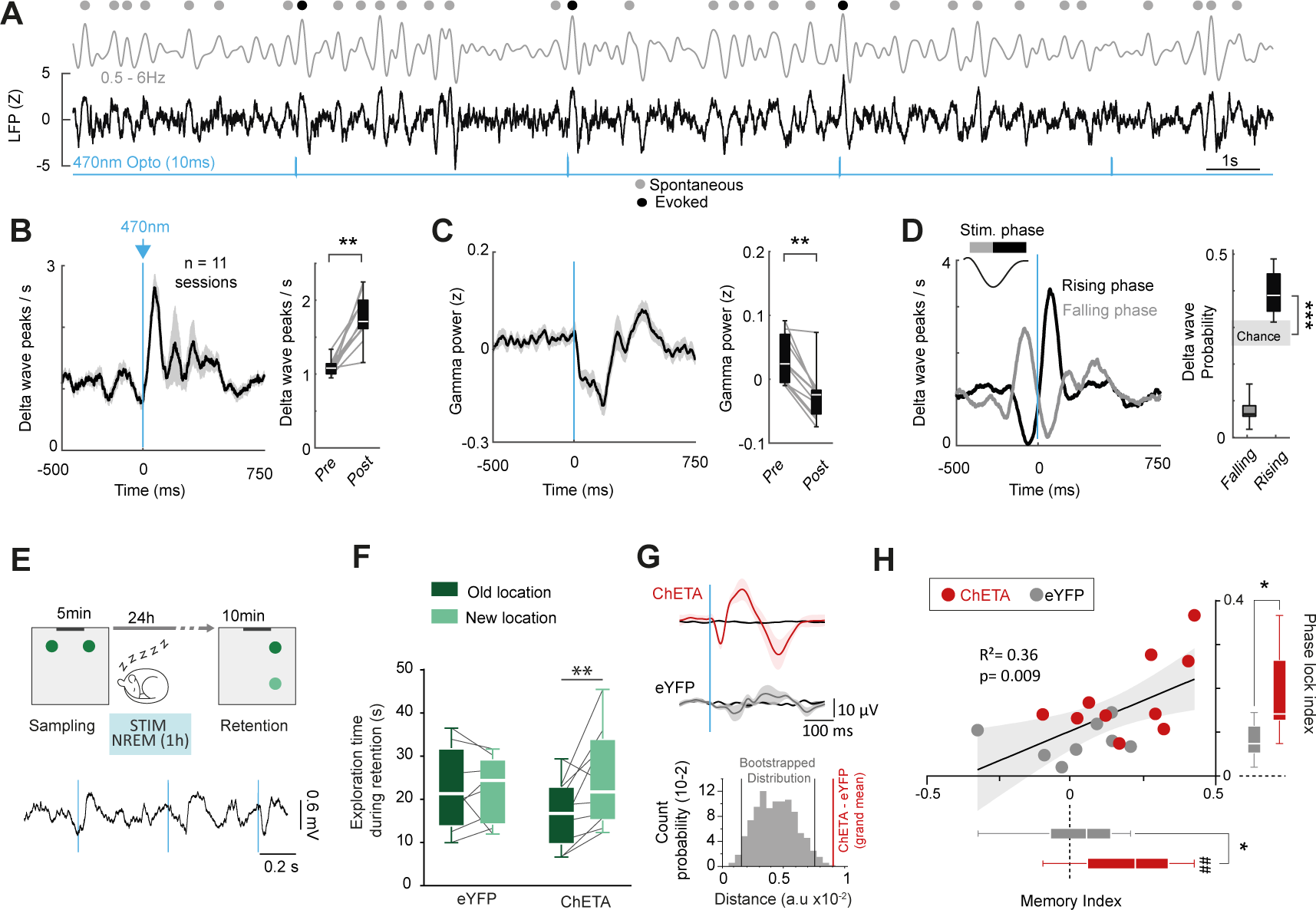
Claustrum stimulation facilitates cortical delta waves and increases memory consolidation. **A**: Field potential recording from the retrosplenial cortex during claustrum stimulation in NREM sleep (raw LFPs - black or filtered for 0.5-6Hz - grey). Evoked and spontaneous delta wave peaks are shown. **B-C**: Claustrum stimulation increases delta wave peaks (B) and decreases gamma frequency power (C). **D**: Claustrum stimulation facilitates the presence of delta waves only when the stimulation arises on the rising phase of depth-negative retrosplenial cortex LFP recordings, but not on the falling phase. **E**: Claustrum stimulation was delivered during NREM sleep following novelty sampling. An example EEG trace from above the retrosplenial cortex during stimulation is below. **F**: The object exploration time during the retention task phase, the day following light delivery into the claustrum. Only the ChETA group exhibited a preference for the new object location (Two-way ANOVA, interaction Object x Virus: F_(1,16)_=4.63, p=0.0470; eYFP: t_(7)_ = 0.04, p = 0.97; ChETA: t_(9)_ = 2.9, p = 0.02). **G**: The mean evoked activity for claustrum stimulation (ChETA, red) and control mice (eYFP, grey) during NREM compared to shuffled stimulation times (black). Dark lines are the mean and shaded lines are standard error of the mean. Below is the bootstrap analysis showing that eYFP and ChETA groups have significantly different responses (see Methods and Materials). **H**: The linear correlation between the memory index and phase locking index. The memory index was significantly greater than zero for ChETA (0.198 ± 0.054, t_(9)_ = 3.61, p = 0.006) but not eYFP mice (0.017±0.06 t_(7)_ = 0.28, p = 0.79), and the ChETA group had an increased memory index relative to eYFP mice (t_(16)_ = 2.22, p = 0.04). The phase locking index was greater in ChETA mice (eYFP: 0.08±0.01; ChETA: 0.18±0.03, t_(16)_ = 2.88, p = 0.01). *p<0.05, **p<0.01, ##p<0.01 (rank sum test against 0).

It is well known that delta frequency and the low cholinergic tone during NREM sleep facilitates memory consolidation, and artificially enhancing synchrony during NREM sleep increases memory performance in rodents and humans ^30,75–77^. It was previously shown that claustrum lesions decrease cortical slow-wave states during NREM sleep ^7^, but it is unclear if the claustrum can participate in sleep-dependent memory consolidation. Therefore, we optogenetically stimulated claustrum neurons during NREM sleep following novelty encoding to determine if we could enhance consolidation of an otherwise labile memory trace. We used a novel object location task, where mice explore one pair of objects, followed 24-hours later, by shifting one of the objects to a new spatial location ^75,78^ (**Figure 4E-H**, **Methods and Materials**). Mice demonstrate their detection of the new location by an increased relative exploration of the displaced object. A single 5-minute sampling period is not sufficient for effective object discrimination 24-hours later ^75^. Indeed, control mice (eYFP in the claustrum) receiving 2Hz claustrum light pulses during the first hour of NREM post-acquisition did not exhibit object discrimination 24-hours later. However, mice receiving claustrum stimulation during post-acquisition NREM sleep showed an increased relative exploration of the displaced object the following day (**Figure 4F, H**), indicating enhanced memory consolidation. In addition, there was a positive correlation between the claustrum-evoked delta phase locking and the memory index, suggesting the efficacy of claustrocortical modulation drives the increase in consolidation (**Figure 4G-H**). Mice receiving claustrum stimulation during the sampling period did not show an increase in the performance on the test-day (**Figure S6**). These results show that claustrum-retrosplenial stimulation during NREM sleep most effectively improves episodic memory consolidation through the phase-entrainment of cortical delta waves. Together with the knowledge that claustrum neurons are most active and most effective at communicating with the cortex during NREM sleep, these behavioral data suggests that the claustrum-retrosplenial circuit is wired for function specifically during “offline” synchronized states.

## Discussion

Our results provide a detailed measurement of the brain-state dependent organization of claustrum neural activity, complimenting other recent work describing claustrum activity in sleep and waking states ^6,7,12,13,16,37,49,79^. We show that claustrum-retrosplenial projections are more active during synchronized cortical states such as NREM sleep and reduced during desynchronized states such as arousal and movement. In addition, claustrocortical communication was the most efficient during synchronized states compared to desynchronized states. Muscarinic cholinergic blockade enhanced claustrocortical signaling suggesting that the reduction in claustrocortical communication during desynchronized states is mediated in part through the actions of acetylcholine. Activating claustrum neurons during NREM sleep temporally aligned cortical delta waves, which led to increased memory consolidation. Collectively, these results suggest that activity within this particular claustrocortical projection stream favors a functional role during ‘offline’ brain activity such as that involved in memory consolidation.

While it has been reported that claustrum cells are more active during NREM than wake ^7^, it has remained unclear to what extent claustrum neurons change their activity as a function of other brain and behavioral states. In rodents, there have been contradictory results regarding the activity of claustrum neurons in REM sleep. It was shown that claustrum-retrosplenial neurons exhibited increased immediate early gene activity (c-fos) during REM sleep rebound ^37,80^. However, fiber photometry of claustrum-cingulate neurons demonstrated neural suppression during REM ^13^. Our imaging approach allows the detection of both increases and decreases in individual axons, which helps reconcile these previous discrepancies. Indeed, a small population of claustrum neurons could express c-fos with persistent REM conditions. However, on average these claustrum-retrosplenial neurons are inhibited during this state. Another report suggests the claustrum homolog in reptiles exhibits alternating patterns of coordination between claustra of each hemisphere during REM ^49^. Our results suggest there could be species differences in claustrum activity, at least within this projection to midline cortex, as it appears as though claustrum activity during REM in mice approaches a non-functional low activity state. However, future work perturbing activity in different brain states (across species) will be required to make a definitive assessment of what function the modulation of claustrum neurons during REM sleep serves.

Previous work has focused predominantly on the measurement and manipulation of claustrum neural activity in tasks involving sensory related decision making known to involve areas of the frontal cortex ^9–11,13,81–83^. Interestingly, activity within the claustrum-anterior cingulate projection has been shown to predict the behavioral performance on a sensory selection task ^13^. Specifically, increased claustrum activity occurred when the animal failed to report the detection of salient sensory stimuli and reduced claustrum activity was associated with impulsive motor action, suggesting a hypervigilance. These results are aligned with our data showing claustrum activity is inversely correlated with the level of arousal and brain-state desynchronization. Another study in rats has shown that a population of cells in the anterior claustrum are negatively correlated with running speed^79^ supporting the concept that that brain and behavioral states associated with strong desynchrony (locomotion) are accompanied by less claustrum activity (**Figure 2**).

The manipulation of claustrum neurons has been shown to impact performance in several tasks ^8,11,12,82,84^. However, the manner in which this change arises has not been established. Neural activity patterns during periods of rest often predict or participate in the shaping of future behavior even in complex cognitive tasks ^6,85–87^. Therefore, it is foreseeable that the impact of claustrum perturbation on behavior arises during ‘offline’ synchronized states that are believed to participate in aspects of consolidation or motor planning. For the most part, brain and behavioral state has not been considered when measuring or manipulating the claustrum in awake animals. Therefore, future work probing claustrum function could benefit from measuring brain state fluctuations (using field potential measurements) to potentially explain what aspect of different behavioral tasks involves the claustrum. It will be important that future behavioral studies employ temporally precise manipulation of claustrum activity to relate any changes in behavior to the brain state at the time of manipulation.

Several questions arise from the results we show here. First, what impact does claustrum activity have on the precise timing of cellular activity in the cortex during NREM or other synchronized cortical states? Previous work suggests the prefrontal cortex undergoes a ‘downstate’ or period of activity suppression during claustrum bursts. However, in the retrosplenial cortex, many more excitatory cells undergo feedforward activation prior to feedforward inhibition ^41^. In addition to delta waves, ripple activity has also been described as a key player of hippocampal-cortical coordination during NREM sleep and memory consolidation ^86,88^. Given the link between claustrum activity and memory consolidation, future work can begin to assess how claustrum neurons may participate in the cellular and network activity patterns known to be important for sleep-dependent memory consolation. One caveat of our study design is that we did not analyze claustrum activity in context of innate or learned sensory-motor behaviors as done previously^6,13–15^. One recent report identified licking-related claustrum neurons projecting to somatosensory cortex, in a sensory detection task and showed that these neurons were robustly modulated by ongoing and future motor action^6^. Further work will therefore be required to determine how separate populations of claustrum outputs are jointly modulated by brain state and sensory-motor variables.

## Acknowledgements

We thank The Faculty of Medicine & Dentistry Cell Imaging Center for support, and Anna Taylor and Jeremy Cohen for comments on the manuscript. Funding was provided by the following: Canada Foundation for Innovation John R. Evans Leaders Fund (JELF), Grant/Award Number: 37931, Canadian Institutes of Health Research, Grant/Award Number: 426485, National Alliance for Research on Schizophrenia and Depression, Natural Sciences and Engineering Research Council of Canada, Grant/Award Number: RGPIN2018-05212, and CNRS International associated laboratory, LIA JOURNEY. JJ holds a Canada Research Chair award.

## Author contribution

Design/Conceptualization: All authors. Experiments and Methodology: BM, CP, AD, RG. Analysis: All authors. Supervision: RG, JJ; Writing: All authors. Funding: RG, JJ.

## Data availability statement

Data are available from the corresponding authors upon request.

## Supplementary Materials

Materials and Methods, Supplementary Figures 1-6

## Methods and Materials

### Animals

All procedures were performed according to the Canadian Council on Animal Care Guidelines and the European Committee Council directive (2016/63/UE) and were approved by the University of Alberta Animal Care and Use Committee or the French Ministry of Research (APAFIS#20388-2019042517013497). Imaging was performed in male and female C57BL/6 mice (60 to 240 days old). Head-fixed optogenetic experiments were conducted in adult male mice (60-240 days old), and behavioral optogenetic experiments were performed on adult male and females CD1 mice (60 to 150 days old). Mice were group-housed in a temperature-controlled environment on a reverse 12-hour light-dark cycle and provided with ad-libitum food and water.

### Surgery

Mice were given 5mg/kg of carprofen via ad-libitum water 24 hours before and 72 hours after surgery. Mice were initially anesthetized in an induction chamber with 4% isoflurane mixed with pure oxygen and then maintained at 1.5-2.0% isoflurane when placed on the stereotaxic frame. An electric pad kept the body temperature at 37°C during the surgery. Mice were administered a local anesthetic (0.5% bupivacaine) subcutaneously before the skin was incised along the midline to expose the skull. The skull was cleaned and kept moist during surgery with 0.9% saline. The head was then levelled in rostral-caudal and medial-lateral directions according to bregma. The procedure for each specific surgery following this step is detailed below.

### Virus injection and brain coordinates

Craniotomies were marked and manually drilled using a 400µm dental drill above the retrosplenial cortex and the claustrum. Anterior-posterior (AP) and mediolateral (ML) coordinates were measured relative to bregma. Dorsal-ventral (DV) coordinates were set from the brain surface. Retrosplenial cortex coordinates were AP: -1.8mm, ML: +/-0.4mm and DV: -0.5mm. Claustrum coordinates were AP: +1.25mm, ML: +/-2.5 to 2.6mm, and DV: -2.5mm. Injections were made by lowering pulled glass micropipettes (10-20µm) filled with mineral oil and loaded with specific adeno-associated viruses (AAV) into the brain. AAVretro-hSyn-Cre (Addgene, Catalog number 105553) (150nl) was injected bilaterally into the RSP. The claustrum was injected with either 200nl of AAV5-hSyn-DIO-EGFP (Addgene, Catalog number 50457), 200nl of AAV1-syn-FLEX-jGCaMP7f-WPRE (Addgene, Catalog number 104492), 200nl of AAV1-syn-FLEX-jGCaMP8-WPRE, or 200nl AAV5-EF1a-DIO-hChR2(H134R)-EYFP (Addgene, Catalog number 20298),or AAV5-EF1a-DIO-ChETA-eYFP (Addgene, Catalog number 26968), depending on the experiment. After each injection, the pipette was allowed to rest for 10min in the brain to prevent backflow. At the end of the surgery, the skin was sutured, and the mice returned to clean heated cages.

## Two photon calcium imaging and analysis

### Skull thinning and head plating

Two to three weeks following virus injection, mice were planted with a headplate, electrodes, and the skull thinned for imaging using general surgical preparation described above. Using a 1mm dental drill bit, the skull over RSP was thinned by laying the drill horizontal and dragging the drill bit across the skull surface using moderate speeds on the variable control. After each thinning pass, saline was added to limit heating. The skull was thinned until flexible and the vasculature could be observed using a dissecting microscope. Using a 33-gauge needle, a small drop of superglue was applied to cover the thinned area, and a custom coverslip (∼3-4mm wide, ∼2-3mm tall) was pressed into the glue over the thinned area and held in place with forceps until the glue set. Craniotomies were then drilled for a stainless-steel support screw (1mm diameter, 8L0X3905201F Protech International), local field potential electrodes, and a reference electrode. Electrodes were made of coated stainless-steel wire (0.008/0.005 in diameter coated and uncoated, respectively, 791100 A-M Systems) soldered to gold male miniature pin connectors (520200, A-M systems). LFP electrodes were implanted as close as possible to the imaging window, usually in secondary visual cortex, and the prefrontal cortex (A/P: +1.7, M/L 0.50, D/V -1.0). Several mice had EMG electrodes implanted in place of the second LFP electrode. The reference electrode was implanted in the cerebellum. Electrodes were secured with a small amount of dental cement (Teets Cold Cure Dental Cement). A custom stainless steel headplate with an oval shaped window was then implanted and a high strength dental cement (C&B Metabond) was used to fill in the remaining area of the skull and headplate window and electrodes. Mice were allowed to recover from surgery for over a week before starting habituation.

### Habituation

Head-plated mice were first habituated to the experimenter for several days before becoming affixed to the headplate holder for the first time. To habituate the mice to the microscope, the amount of time head-fixed to the microscope increased over consecutive days of recordings. Several mice were observed to enter REM sleep during the third day. Subsequent imaging sessions were performed across 1-4 months following head-plating. Only mice that had visible axon fibers and usable LFP recordings were used for imaging. Following sleep recordings, mice were then trained on the treadmill for locomotion recordings. Mice were habituated on the treadmill for 1-2 days before continuing imaging. Following sufficient treadmill recordings, calcium imaging was recorded following either saline (0.9%) or atropine sulfate (10 mg/kg) injections.

### In vivo imaging

An upright ThorLabs Bergamo II microscope with a Ti:Sapphire femtosecond laser (Thorlabs Tiberius) and 8kHz resonant scanner (Thorlabs) was used for in vivo imaging, with the laser set to 920nm for jGCaAMP8m imaging. Two-photon emission was filtered using a 562nm long pass dichroic mirror (FF562-Di03-32×44, Semrock) and 525nm emission filter (ff03-525/50-32, Semrock) before entering a GaAsP PMT (PMT2100, Thorlabs). The main microscope body was inside a light tight enclosure with sound dampening foam (5692T49, McMaster Carr) on all sides, roof, and door. Head-fixed mice were either attached to an enrichment tube holder cut in half lengthwise for the initial habituation and sleep recordings or manual treadmill(*42*) with a rotary encoder to track movement for locomotion recordings. Notches were cut in the tube to allow the headplate holders to be placed at a level that would keep the mice at a comfortable height. Square white nesting pads torn in half were attached to the bottom of the tube to provide cushioning. A Nikon 16x 0.8NA objective was used for 512×512px imaging at 30 frames / second. Bidirectional correction was adjusted per field of view based on location and depth of imaging. Acquisition settings were 4x zoom, 210.36×210.36µm field of view, 0.411um/pixel. Laser power was set to 30-150mW at the objective depending on skull thinning and GCaMP expression and depth of imaging. Laser power was modulated through a quarter wave plate (Thorlabs Power Modulator) to achieve a sufficient signal to noise above the noise floor of the detectors.

### Pupil and face motion

To capture pupil and face motion, an IR sensitive CS-mount USB camera (FMVU-03MTM-CS, FLIR) with a 15-50mm f1.7 lens was used. Imaging was performed at 30Hz synchronized with calcium imaging frames. To decrease the minimum focusing distance and enable greater macro magnification, a C mount adapter was attached between the camera and CS mount lens. A DC infrared LED array was powered via 12V battery pack inside the microscope box and was positioned to illuminate the pupil. Pupil size was calculated on a cropped version of the full-face movie. Each image in the movie was inverted and thresholded to highlight the pupil relative to the background. The threshold was determined for each mouse and imaging session. The image subregion containing the pupil was filtered using a two-pixel median filter and an ellipse fit to the pupil. The area of the ellipse was taken as a relative measure of pupil size in each imaging session. The data were z-scored for analysis. Imaging frames where the mouse closed its eye or where a pupil could not be detected were interpolated using adjacent frames. Pupil phase was calculated by z-scoring and filtering the pupil size using low pass filter (0.1Hz) and then taking the angle of Hilbert transformed signal. Pupil dilation was defined as the rising phase from -120 degrees to 0 degrees where -180 and 180 degrees signify the trough, and 0 degree is the peak. For the analysis of facial motion, a region of interest was placed on the vibrissae pad. The absolute value of the mean difference in pixel intensity between adjacent frames was calculated for all adjacent frames(*49*, *51*) and severed as a measure of facial movements.

### Treadmill locomotion

Treadmill speed was recorded using Arduino that converted the distance moved on the treadmill into a voltage that was linearly related to speed. Locomotion was down sampled to the same frequency as calcium imaging by averaging the treadmill speed across 60ms centered on the onset of the calcium imaging frame. This vector was smoothed with a 2s moving average, and periods of movement greater than 1cm/s were used for analysis. Brief periods of movement lasting less than 2s were not considered locomotion epochs. Periods of movement separated by less than 2s were considered part of the same movement epoch.

### Local field potentials (LFPs)

LFP and reference electrodes were connected to a head stage (A-M Systems Model 1700) and connected to a differential amplifier (A-M Systems, Model 1700 Differential AC Amplifier). Signals were high pass filtered at 0.1 or 1Hz and lowpass filtered below 1kHz and digitized at 30kHz on the Thorlabs system. LFPs were synchronized with calcium imaging using a binary pulse that was sent to the digitizer for every imaging frame. LFPs were analysed using the chronux signalling processing toolbox(*82*). Power spectrums were calculated on three second periods of data centered on each imaging frame, using the multitaper spectrogram function in chronux. Therefore, each imaging frame had an associated spectrum. The mean delta power (0.5-4Hz), theta (6-11Hz), and gamma (30-100Hz) power were extracted, and the theta-delta and gamma-delta ratios used for analysis.

### Analysis of calcium imaging data

Imaging movies were motion corrected in the xy plane using either MOCO(*83*), or Suite2P(*84*) (both gave comparable results). Movies with clear z-plane drift were not analyzed (these typically occurred early in habituation when mice were getting accustomed to the head-fixation). Claustrum axon regions of interest (ROI) were manually defined based on the maximum intensity projection or standard deviation of the imaging session. ROI were made to specifically avoid sampling the same axon more than once in an imaging session. To objectively remove instances where the same axon was contained as part of multiple ROIs we calculated the correlation coefficient between all pairs of ROI, and in cases where pairs of ROIs had a correlation greater than 0.7, the ROI with the greatest overall change in fluorescence (ΔF/F) was saved and the other ROI removed from analysis. ΔF/F for each ROI was calculated by subtracting and dividing the raw trace by the lower 40th percentile of a fifteen second moving window. ΔF/F traces were deconvolved using the OASIS toolbox(*85*). From the deconvolved data, we measured the number of individual events (ignoring absolute amplitude) which was converted to an event rate (events/s). Unless otherwise indicated, we used either the deconvolved ΔF/F calcium traces or the calcium event rate (a more conservative estimate of activity). The population claustrum activity was defined as the mean event rate or deconvolved calcium signal, averaged across all ROI as a function of time. Between imaging sessions, a new group of ROI were selected, and we did not attempt to track axons across days, so it is foreseeable that some ROI were inadvertently sampled on multiple sessions. For this reason, our conclusions were arrived at through an analysis of single ROI, the mean claustrum population activity, and the averaged population activity within and across animals. All major claims were supported at all levels of analysis (ROI, sessions, mice).

Control imaging with GFP labelled claustrum axons confirmed no observable z-drift on the imaging setup during wake and sleep (**Figure S1**). For locomotion, we used auto fluorescence blebs that showed higher background fluorescence without observable calcium transients(*47*, *51*), and measured their change in fluorescence across time, in order to ensure a lack of z-motion (**Figure S2**).

The correlation between claustrum ROI (or claustrum population activity) and other variables was performed using the ‘corr’ function in Matlab. The time-lag correlation analysis was performed by calculating the correlation between two variables, shifting the one variable by different positive or negative time lags. The correlation asymmetry was calculated by taking the mean time-lag correlation coefficient in the from the positive time lags ranging from 0-5s minus the mean correlation coefficient within the -5s -0s time lag interval.

### Head-fixed sleep-wake scoring

Sleep-wake scoring in head-fixed mice was performed using LFPs, pupil, and face motion and was carried out similar to previous reports(*43*–*45*). We used an automated sleep-wake scoring procedure. The theta (6-11Hz) to delta (0.5-4Hz) spectral power ratio was calculated on 3s epochs centered on each imaging frame to obtain a theta-delta ratio value as a function of calcium imaging data. The theta-delta ratio, pupil, and face motion data were each smoothed using a 10s sliding window, and z-scored. NREM was defined as periods where pupil and face motion z-score were less than 0 and the z-score of the theta-delta ratio less than 0. REM sleep was defined as periods of time when the z-scored pupil was less than 0, and z-scored theta-delta power greater than 2 standard deviations above the baseline theta-delta ratio. Baseline theta-delta was calculated using the lowest 95^th^ percentile of the data within a session. Rapid eye movements during the detected REM periods provided further evidence that the mouse was in REM sleep. All other periods were defined as quiet wakefulness. REM epochs lasting less than 15s and flanked by NREM were re-scored as NREM. REM epochs separated by less than 20s were interpolated as part of the same REM epoch. Likewise, periods of NREM less than 5s flanked by wake were rescored as wake. REM epochs were required to last at least 30s to be included in the analysis otherwise these periods were removed from analysis. Wake periods less than 5s flanked by NREM periods were rescored as NREM. A total of 72.2 minutes of REM were detected with individual epochs lasting 2.1±0.7 minutes. This sleep-wake scoring procedure is applicable only for data sets with known sleep-wake transitions between different states (usually lasting >15 minutes) in head-fixed mice.

## Head-fixed optogenetic experiments and analysis

### Surgery

Mice were injected with AAVretro-hSyn-Cre in the retrosplenial cortex and AAV5-EF1a-DIO-hChR2 in the claustrum using surgical procedures described above. Two weeks later, 200µm fibre optic cannulas (RWD, Catalog number R-FOC-BL200C-39NA) were positioned, bilaterally, above the claustrum (-2.1mm ventral to brain surface). An opaque adhesive cement (C&B-Metabond, Catalog number S380) was applied around the ferrule of the optic fibres. LFP electrodes were implanted into deep layers of the retrosplenial cortex (A/P: -2.0mm and ML: 0.4mm, D/V: - 0.5) and prefrontal cortex (A/P: +1.8mm and M/L: 0.4mm, D/V: -0.8). A reference electrode was placed into the left side of the cerebellum. A bone screw was inserted into the skull on each end of the head plate, one above the left prefrontal area and one around the visual cortex area. Adhesive cement covered all remaining skull areas, the screws, and the head plate.

### Head-fixed awake optogenetic experiments

For awake experiments, mice were head-fixed and allowed to run on a linear treadmill in a dimly lit (40 lux) environment. Mice were habituated to the treadmill for seven days. During recording sessions, 10ms 470nm pulses (Laserglow Technologies) were bilaterally delivered to the claustrum at 0.5Hz or 2Hz using a Master 8 controller (A.M.P.I) while mice spontaneously transitioned between locomotion and rest. Before the experiment, the laser power was adjusted (2-7mW) for each mouse so that the maximum change in LFP amplitude did not exceed the dynamic range of spontaneously occurring LFP signals. The running speed of the mouse was quantified using a treadmill as described above. Electrodes were connected to a differential AC amplifier (A-M Systems, Model 1700) for amplifying the RSP local field potential signal of ×10k. The signal was filtered between 1Hz and 1000Hz at the amplification stage and acquired at 1219Hz. A camera recorded pupil size and face movement as described above. Each mouse underwent 5-7 stimulation sessions separated by at least 48 hours.

### In vivo pharmacology

Pharmacological experiments were all performed on the head-fixed mice. The baseline period consisted of a 20-minute recording where mice spontaneously transitioned between locomotion and rest. Mice were then briefly anesthetized with isoflurane before receiving a dose of either atropine to block muscarinic receptors (10mg/kg), physostigmine, an inhibitor of acetylcholinesterase (0.6mg/kg) or saline (0.9% sodium chloride, subcutaneously). Post-injection data was recorded for 60-80 minutes. Throughout pre- and post-drug or saline recordings, the claustrum was continuously stimulated with 10-ms pulses of 470nm light at 2Hz as detailed above in blocks of 30s separated by 5 minutes.

### Head-fixed sleep optogenetic experiments

Mice were habituated to the sleep tube as described above. During recording sessions, 10ms 470nm pulses were bilaterally delivered to the claustrum at 0.2Hz during wakefulness and NREM and 0.5Hz during REM. Electrodes were connected to a D/C coupled pre-amplifier head stage and amplified and digitized as described above. The retrosplenial cortex electrode, the timing of light stimuli and the pupil size were acquired synchronously for post-experiment analysis. Each mouse underwent 2-5 sleep stimulation sessions and results were averaged across sessions for each mouse.

### LFP analysis

The putative monosynaptic post-synaptic potential was measured by first averaging the LFP response across all trials within a defined brain or behavioral state. This average trace was smoothed (5ms sliding window), and the absolute slope and amplitude were calculated on this final LFP waveform from 10-100ms following the start of the light pulse for each state. To compare across states measured during sleep experiments and experiments on the treadmill, we normalized the evoked response to the quiet wakefulness state, which was obtained in both awake and sleep experiments.

### Delta wave detection

Delta waves were detected as described previously (*71*), where field potential data were filtered between 0.5-6Hz, and zero-crossings of the first derivative of this signal were detected corresponding to the start, peak, and end of each wave. Delta waves were classified as waves with a peak >2 or peak >1.5 and end trough <1.0. This led to ∼ 28% of the potential oscillation cycles within NREM sleep being classified as a delta wave. Gamma frequency power was measured by taking the z-score of the absolute value of the Hilbert transformed gamma (30-100Hz) amplitude as a function of time. The phase of optogenetic stimulation was determined using the angle of the Hilbert transform of the filtered data.

## Novel object location task and analysis

### Novel object location task

The novel object location (NOL) task used in this study is based on mice’s preference for novelty. This task was performed in an open field (square or hexagonal depending on the condition) with a spatial cue fixed on one of the walls. The device was illuminated by indirect LED light (open field center: 30 lux) and a radio played a background noise (open field center: 45 dB). Before the task, all mice received a habituation trial of 10 min once a day for 3 days, with a different object placed in the center of the open field at day 2 and 3. For the NOL task, mice first explore the open field with two new and identical objects placed in the corner at equal distance from the spatial cue during a 5 min acquisition phase. 24 hours later, they performed a 10 min retention test where they explore the same open field, in the presence of the same two identical objects, only the spatial configuration was changed: the less explored object during the acquisition phase was displaced in a new location (in the empty corner adjacent of its initial location). During each phase of this task, the mice were connected to electrophysiological recording and optogenetic stimulation devices. Between each trial, the walls, floor, and objects were wiped with 35% ethanol.

### NOL analysis

Object exploration time was recorded and defined for each object as the nose pointing toward the object within 2 cm. A memory index was calculated and defined as: (time spent exploring the displaced object − time spent exploring the non-displaced object)/total exploration time for the two objects during the 10-min retention test. This task was performed for two conditions depending on the period of optogenetic stimulation. In the sampling condition, the mice were optogenetically stimulated at 2Hz during the 5-min acquisition phase and in the NREM condition, they were stimulated at 2Hz in their home cage during the first hour of NREM sleep following the acquisition phase. If the mouse entered REM, the stimulation was terminated. All the mice did first the NREM condition and then the sampling condition. For each condition, a different open field was used and randomly assigned to one task: one square open field (50×50×50 cm) with Plexiglas black walls and one hexagonal open field with Plexiglas grey walls (6 sides of 23 cm, 40 cm height). For each test, a unique pair of objects was used, different from one condition to another.

### Freely moving EEG analysis

EEG was recorded through a small screw placed above the retrosplenial cortex and was sampled at 2KHz (Intan amplifier, IntanTech, California, USA) throughout the sampling and the test phase of the NOL Task but also during the 2 to 3 hours immediately following the sampling phase. For the NREM condition, EEG was used to detect NREM sleep and start the optogenetic stimulation. To determine the mean cortical response to claustrum stimulation, EEG traces were first low pass filtered at 30 Hz, aligned to the stimulation, and averaged across all the stimulations. To quantify the efficacy of the stimulation, we used the Kuramoto Index (*86*) which provides a measure of phase consistency across trials for each time point relative to the optogenetic stimulation. We first calculated the Hilbert transform on the filtered EEG trace and, as with the mean response, segmented it into windows around each stimulation (-0.1s and +0.4s). For each point in the EEG signal in this window, the Kuramoto index is calculated according to the following formula:

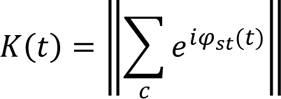

where K(t) represents, for each point *t* of the signal, the absolute value of the sum of the exponential of the imaginary component of the Hilbert transform (ίφ: the signal phase) of the EEG signal at each stimulation (st). Thus, this index indicates at each time t, the phase difference of the oscillations (bounded between 0 and 1). For each mouse and each condition, we used the value of the maximum point of this index in the post-stimulation time period and assigned this as the phase locking index.

To compare and determine the significance of the mean response between eYFP and ChETA mice, we compared the difference between the grand mean response in the two groups to a distribution of bootstrapped datasets that were resampled from the eighteen mice (n=8 eYFP, n=10 ChETA). The absolute value of the difference between two groups were compared. For each bootstrapped iteration (n=1000), the grand mean response for two surrogate data sets were calculated. Each surrogate set contained the mean response from ten randomly selected mice (5 eYFP mice and 5 ChETA mice). Therefore, each of the two groups used in each bootstrapped iteration were comprised of equal number of eYFP and ChETA mice. The difference between group means of the original groups was required to be greater than the 95% confidence interval of the full bootstrapped distribution.

## Histology

Mice were anesthetized with an intraperitoneal injection of a lethal dose of urethane. When unresponsive, mice were transcardially perfused with 1X phosphate-buffered saline (PBS) followed by cold 4% paraformaldehyde (PFA) in PBS. Brains were stored in PFA for a 24h post-fixation at 4°C before being transferred in 1X PBS. Brains were sectioned at 50µm using a vibratome and then mounted with a mounting medium with DAPI (Abcam, Catalog number ab104139). Slices were imaged using a whole slice scanner (Zeiss Axioscan Z1, University of Alberta Cell Imaging Core) to check for cannula placement, electrode location and the expression of channelrhodopsin across different sections of the brain.

## Statistics and Reporting

No statistical methods were used to pre-determine sample sizes. Statistics are indicated in the Figure legends. Unpaired or paired t-tests were used and corrected for multiple comparisons using the Bonferroni-Holm correction. Matlab was used for all analyses unless otherwise indicated. When box plots are used, they show the median, interquartile range, and extreme values of a distribution. When p-values are indicated in Figures, *p<0.05, **p<0.01, ***p<0.001, otherwise the absolute p-value is reported. Figures were constructed using Inkscape or Adobe Illustrator.

---

**Figure S1:**
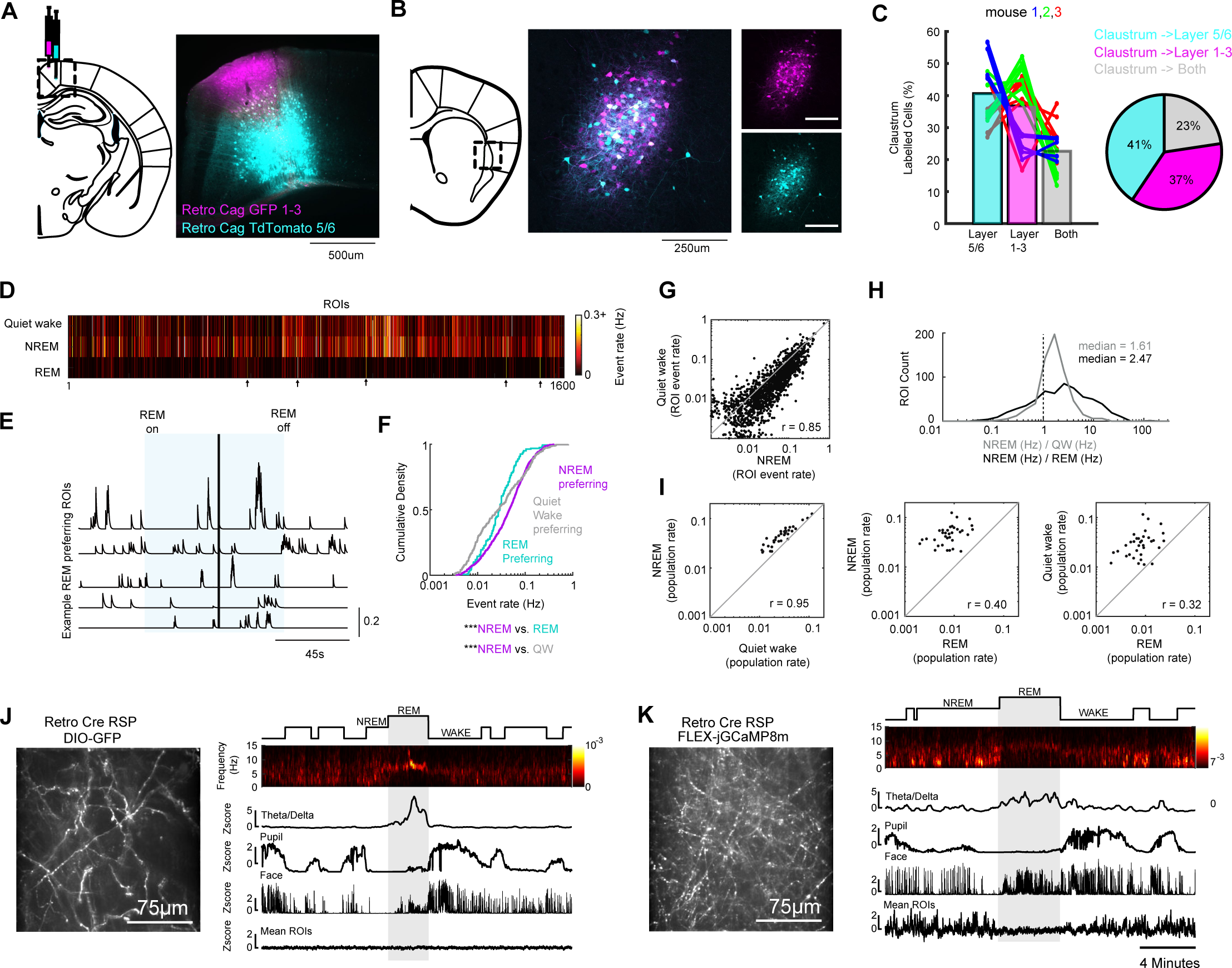
Further analysis of the anatomy and physiology of claustrum ROI in the retrosplenial cortex during sleep-wake states, related to Figures 1 and 2. **A**: Example schematic and image showing virus injection location in the deep and upper layers of the retrosplenial cortex. **B**: Retrograde labelling in the anterior claustrum from the mouse shown in panel A. **C**: The percentage of claustrum cells that were found to project to deep (cyan), upper layers (magenta), or both (grey) for all slices from three mice. There is an upper bound of ∼ 50% colocalization when injecting both tracers into the same region within the retrosplenial cortex (see Marriott et al. 2021). **D**: Heatmap showing the mean Ca^2+^ event rate for all ROIs that were detected in all imaging sessions on the sleep-tube, during different brain states. Arrows below show five ROIs with higher event activity during REM than NREM or wake; these ROIs are the exception as most ROIs were silenced during REM. **E**: The time-series from these REM-preferring ROIs showing the extent of their activation during REM sleep. Note these ROIs are from different imaging sessions, and data are aligned to REM onset and offset as shown above. The vertical line indicates a gap in time where data were spliced together for visualization. **F**: The mean Ca^2+^ event rate for ROIs preferring REM, NREM or quiet wake. The mean event rate for REM preferring cells is significantly less than the mean event rate for NREM preferring cells, suggesting that REM preferring ROIs are likely not a subset of highly active neurons. **G**: The scatter plot of Ca^2+^ event rate of all ROIs that exhibited events in both NREM and quiet wake. A strong linearly relationship was identified. **H**: The event rate ratio between NREM/REM and NREM/quiet wake, across all ROIs that were detected to be active in both states (note ROIs that were silenced in one state are not included here). **I**: The mean population event rate (averaged across ROIs within each imaging session) is shown to visualize the difference between NREM, wake, and REM. The grey line reflects equality between two variables. Correlation coefficients are indicated below. **J**: Control fluorophore imaging does not show changes in fluorescence during REM sleep. Left: an example field of view showing axons in the retrosplenial cortex after cre-dependent GFP was injected into the claustrum instead of GCaMP8m. Axon ROIs were processed the same as the GCaMP data. Right: an example experiment showing there is no considerable change in the average ROI fluorescence across time during REM sleep. **J**: An example GCaMP experiment where the mean z-scored ROI fluorescence decreases during REM.

**Figure S2:**
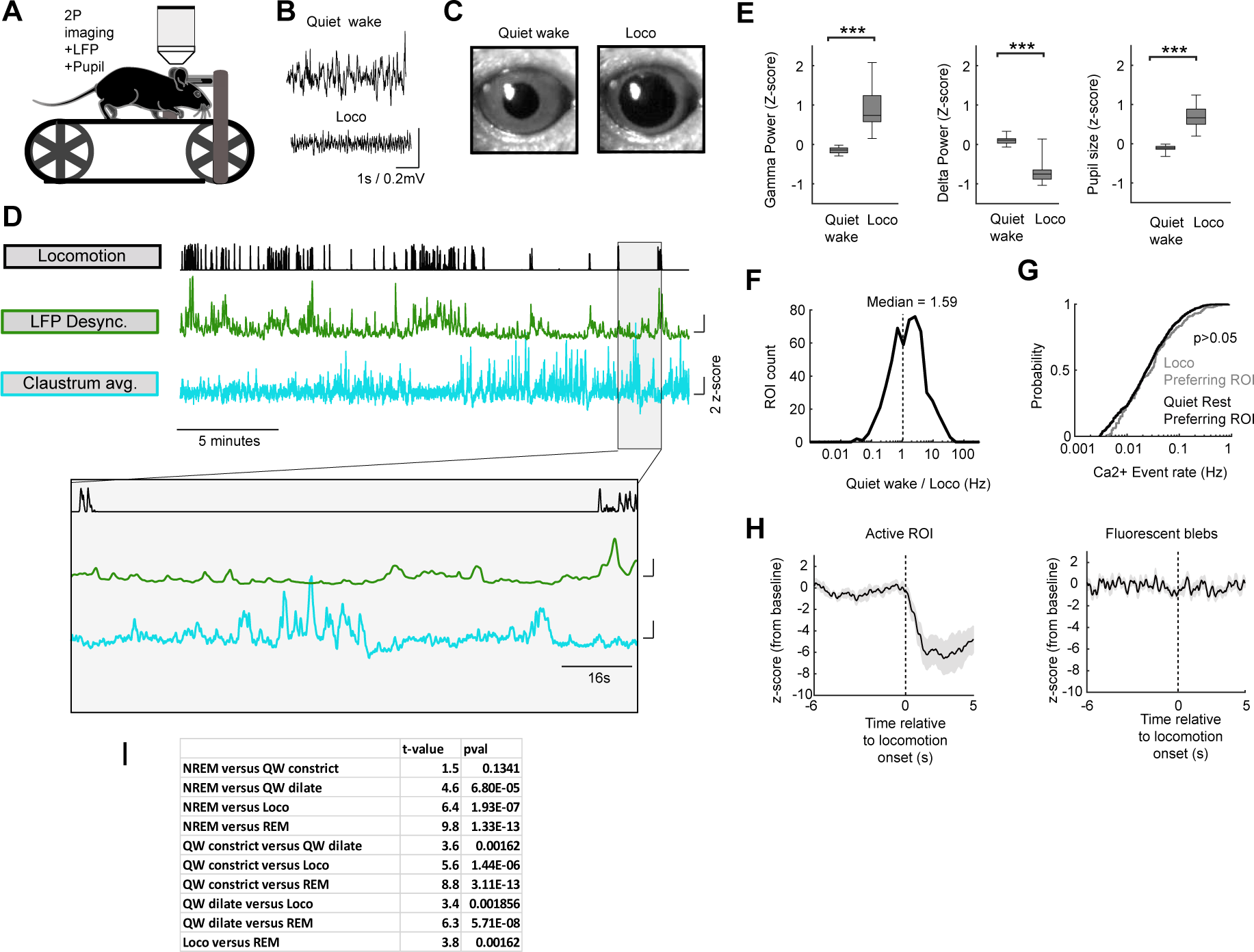
Further information regarding reduced claustrum activity during locomotion, related to Figures 1 and 2. **A**: Schematic showing two-photon imaging setup on the mouse treadmill. **B**: Example local field potential traces during quiet wake and during locomotion. **C**: Example single frames showing pupil dilation during locomotion relative to quiet wake. **D**: An example experiment showing the mouse undergoing rest-locomotion periods together with cortical desynchronization and average claustrum population activity. The shaded area is expanded below. Note the claustrum is mainly active in the 2^nd^ half of the experiment when arousal/locomotion levels decrease. **E**: The group data for LFP gamma power (t_(16)_ = 6.4, p = 9×10^-6^), delta power (t_(6)_ = 7.8, p = 7×10^-7^), and pupil size (t_(21)_ = 12.6, p = 3×10^-11^) for treadmill sessions. Five sessions were removed from analysis due to local field potential artifacts during locomotion. **F**: The ratio of calcium event rates for all claustrum axon ROIs that were active in both quiet wake and locomotion (ROI that were silent during locomotion were not used in this calculation). **G**: The cumulative density function showing that the calcium event rate was not statistically different between ROIs found to be locomotion preferring relative to quiet wake preferring. **H**: Left: The grand mean z-score fluorescence for all ROIs, averaged across session and mouse around locomotion onset. The black line is the mean and shaded area is the standard error from six mice. Right: Neighboring non active autofluorescent blebs in the same field of view were subjected to the same analysis and did not show considerable changes at the onset of locomotion suggesting decreased GCaMP (left) fluorescence is not due to a change in the field of view upon locomotion. **I**: A statistical table showing the pair-wise comparison between claustrum population activity rate across five states, and p-values corrected for multiple comparisons.

**Figure S3:**
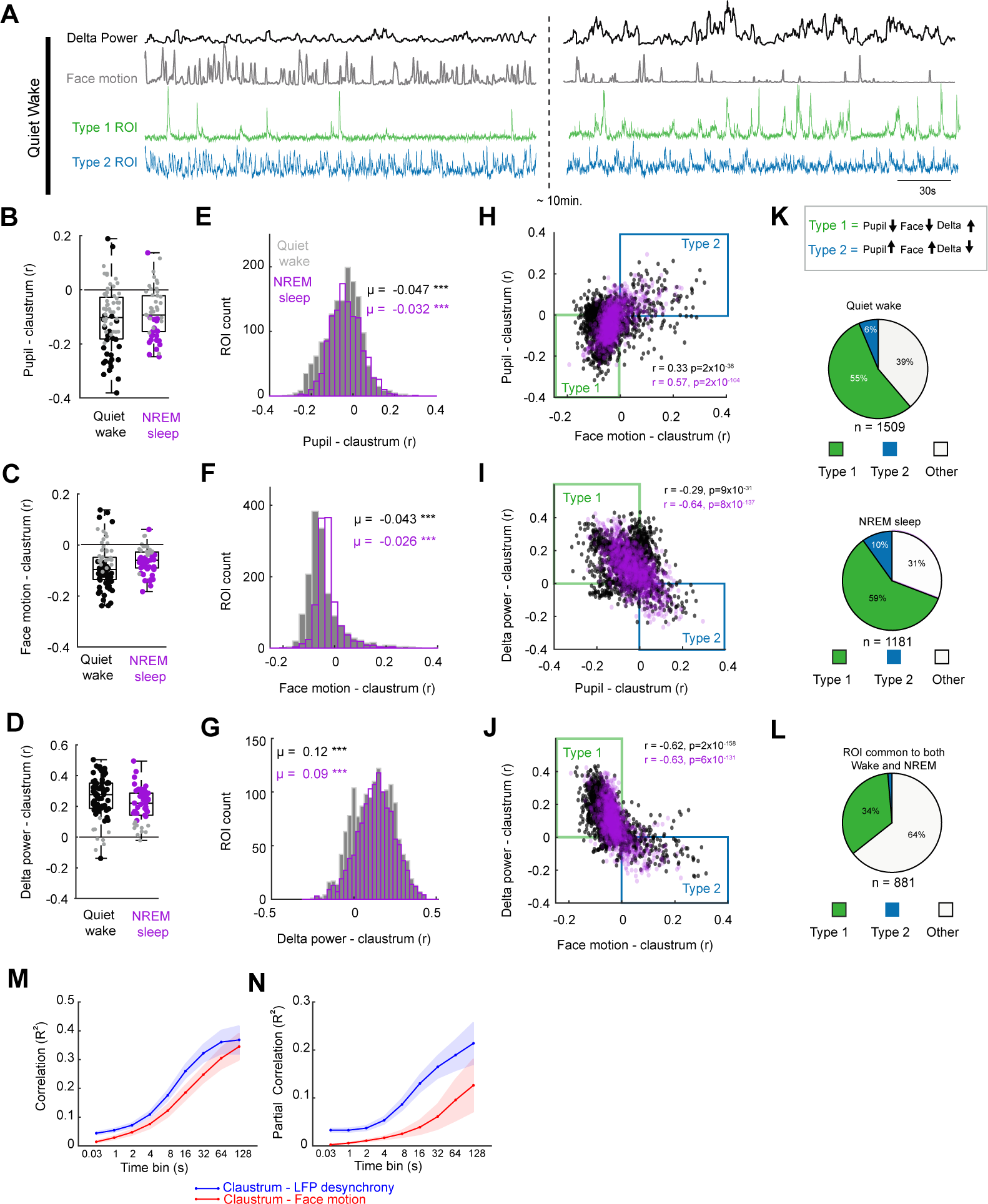
Further analysis and comparison of individual claustrum axon ROI during quiet wake and NREM sleep, related to Figures 1 and 2. **A**: Example ROIs showing Type 1 and Type 2 responses to brain state and behavior changes (see panel K for definitions). **B**: Pearson correlation coefficient between pupil size and claustrum population activity for all sessions for quiet wake (black) and NREM sleep (magenta). Statistically significant sessions are highlighted in black and magenta and nonsignificant sessions are grey. **C**: The same as B, for correlations between face motion and claustrum population activity. **D**: The same as B-C, for correlations between delta frequency power and claustrum population activity. **E**: The correlation coefficient for all claustrum axon ROI with pupil size for quiet wake and NREM. Both distributions where significantly shifted less than 0. **F**: The same as E for ROI correlations with face motion. **G**: The same as **E-F** for ROI correlations with delta frequency power. **H**: The relationship between claustrum-face correlation and claustrum-pupil correlation for all ROIs in quiet wake and NREM sleep. **I**: The relationship between claustrum-pupil and claustrum-delta power correlations. **J**: The relationship between claustrum-face motion correlations and claustrum-delta correlations. **K**: ROIs with a Type 1 activity were those that had a negative claustrum-pupil correlation, negative claustrum-face correlation, and a positive claustrum-delta correlation (green quadrant in panels H-J). ROIs with Type 2 activity were defined as those that had a positive claustrum-pupil correlation, positive claustrum-face motion correlation, and a negative claustrum-delta correlation (Cyan quadrant in panels H-J). Pie charts show the percentage of ROIs detected during quiet wake (top) or NREM sleep (below) for each category. **L**: The proportion of ROIs that maintained a Type 1 or Type 2 activity across NREM and quiet wake. **M**: R^2^ values for claustrum population activity versus LFP desynchrony (blue) and facial motion (red) for different time bins. Correlation coefficients were calculated using the integrated activity over different time bins. The solid lines are the mean and shaded area indicate the standard error of the mean for six mice. **N**: The same as A, using the partial correlation between claustrum population activity and LFP desynchrony or face motion, taking into consideration the other variable.

**Figure S4:**
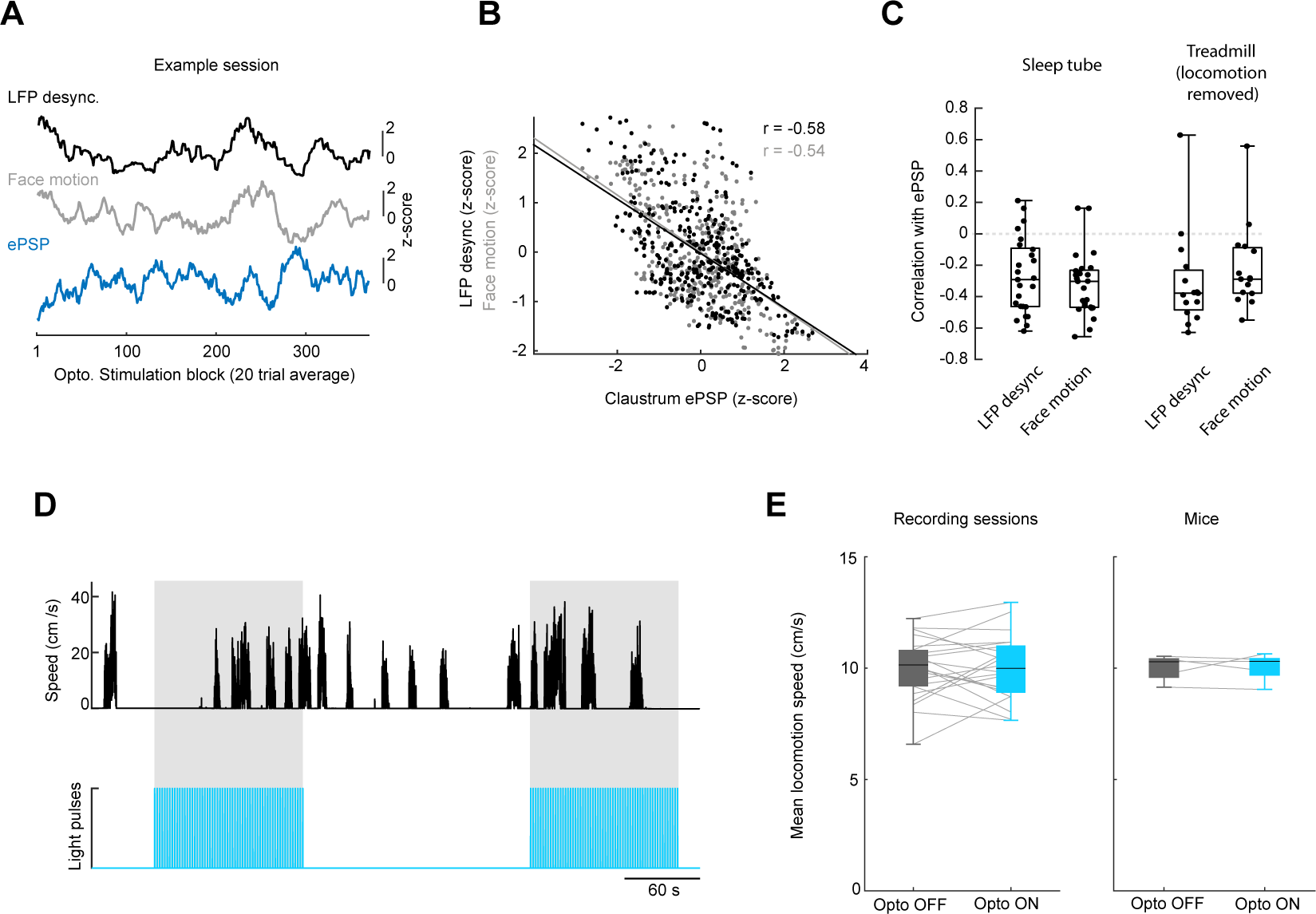
The claustrum evoked cortical response is dependent on brain and behavioral state but does not modify behavioral state. **A**: An example experiment showing changes in the claustrum evoked ePSP size in the retrosplenial cortex as a function of cortical desynchrony and face motion. The ePSP, face motion, and desynchrony were measured in blocks of 20 trials. **B**: Scatter plot of the data in **A** showing a negative correlation between ePSP and both face motion and cortical desynchrony. **C**: The correlation coefficient for each session on the sleep tube and on the treadmill (after locomotion was removed). Each point represents an experimental session. **D**: An example experiment showing blocks of 2Hz claustrum – retrosplenial neuron activation (light blue) and locomotion speed (black). Activation does not interfere with locomotion speed or the ability to initiate locomotion. **E**: Quantification of locomotion speed in the absence or presence of optogenetic activation, showing speed is not significantly altered across sessions or mice.

**Figure S5:**
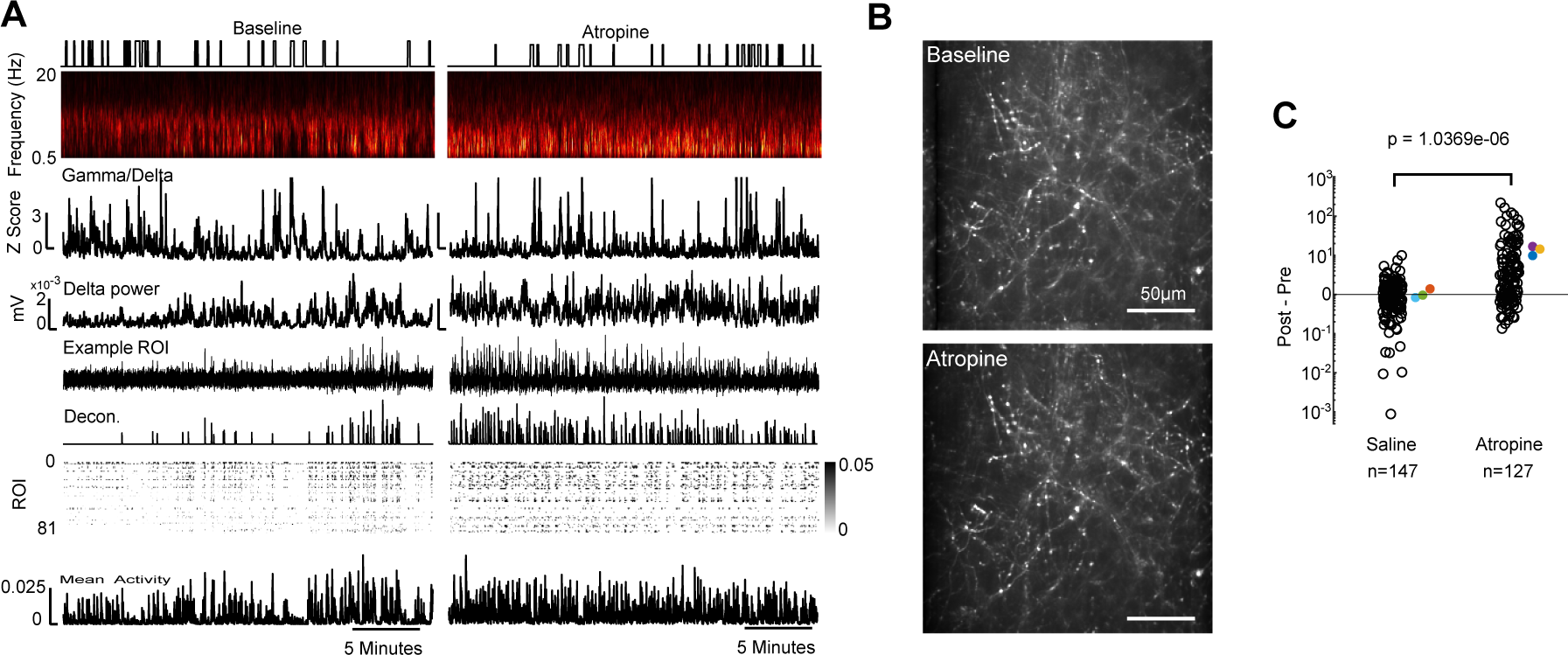
Cholinergic muscarinic receptor blockade increases the activity of claustrum-retrosplenial axons, related to Figure 3. **A**: An example experiment where a mouse was administered 10mg/kg atropine sulfate while performing claustrum axon imaging. The time-frequency spectrogram, LFP desynchrony, delta power, an example ROI (and spike estimation deconvolution shown below), all ROIs, and the averaged claustrum activity. Periods of locomotion are indicated above the spectrogram. Note that atropine increases delta power and claustrum ROI activity. **B**: An imaging field of view pre/post drug. **C**: The mean change in the deconvolved calcium signal for each ROI in saline control (left) or atropine (right) experiments. Only ROIs that were detected in both post and pre sessions were used for these analyses. The colored points indicate the mean data within each of three different sessions (t_(272)_ = 4.99, p = 1×10^-6^).

**Figure S6:**
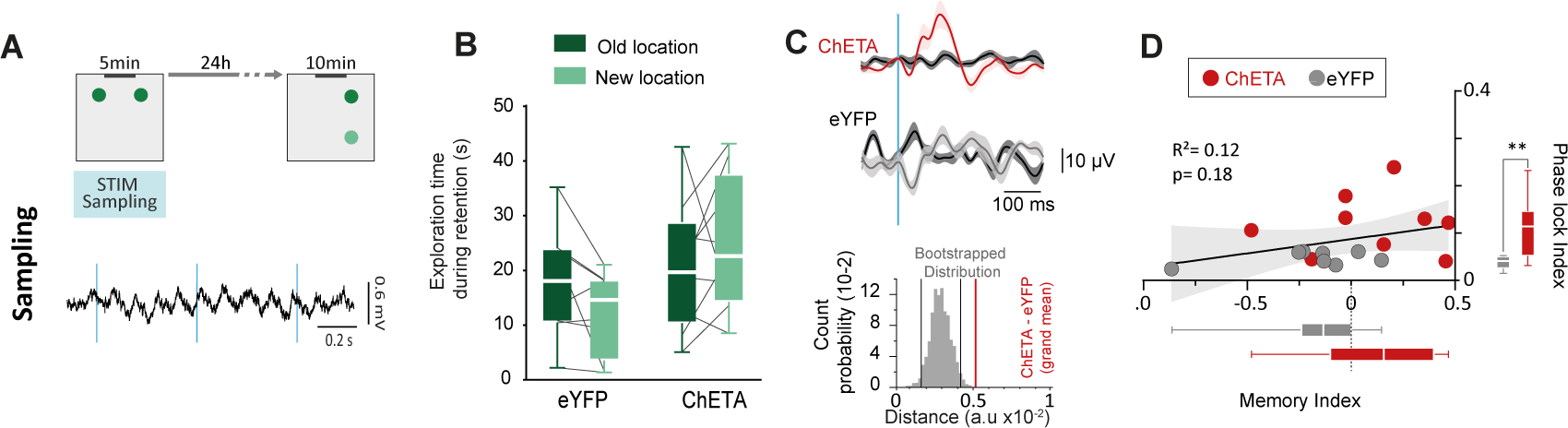
Optogenetic stimulation of the claustrum during sampling phase does not increase memory consolidation, related to Figure 4. **A**: Claustrum stimulation was delivered during the sampling phase of the task while mice explored two objects. Example EEG trace during stimulation during exploration are below. **B**: The object exploration times (in seconds) in the retention task phase, the day following light delivery (2Hz pulses) into the claustrum during the object sampling phase. Neither group showed a preference for the new object location during the retention phase (eYFP: p = 0.06; ChETA: p = 0.45). **C**: The mean evoked activity for claustrum stimulation (red) and control mice (grey) during the sampling phase compared to shuffled stimulation times (black). Dark lines are the mean and shaded lines are standard error of the mean. **D**: The memory index and phase locking index (measured during sampling) were not correlated. The memory index was not different than zero for ChETA (0.102 ± 0.105, t_(8)_ = 0.97, p = 0.36 versus 0) or eYFP mice (-0.18±0.11 t_(7)_ = 1.76, p = 0.12 rank sum test versus 0). The memory index in ChETA mice and eYFP mice was not significantly different (t_(15)_ = 1.93, p = 0.073). However, the phase locking index was greater in ChETA mice (eYFP: 0.04±0.01; ChETA: 0.12±0.02, t_(15)_ = 3.12, p = 0.007). *p<0.05, **p<0.01, ##p<0.01 (rank sum test against 0).

